## Supplementary material for "Individually unique, fixed stripe configurations of *Octopus chierchiae* allow for photoidentification in long-term studies": S1 File. OLAC Ethics Statement

University of California Berkeley
Office of Laboratory Animal Care
203 Northwest Animal Facility
Berkeley, CA 94720-7150
www.olac.berkeley.edu

December 22, 2021

To whom it may concern:

The Roy Caldwell lab from the department of Integrative Biology has performed a study regarding the individually unique, fixed body patterns of Octopus chierchiae. This study was performed at the University of California, Berkeley in an AAALAC accredited facility that is overseen by the Office of Laboratory Animal Care. Since the species is an invertebrate, it is not covered by the USDA or PHS policy. However, because the facility is AAALAC accredited, the building the octopus was housed in is managed and maintains the standards set forth by the Guide for the Care and Use of Laboratory Animals. Additionally, this study did not require an IACUC approved Animal Use Protocol.

UC Berkeley performs all studies ethically and always considers the health of the animal with utmost importance.

Gregory W. Lawson DVM

Gregory W. Lawson DVM, PhD, DACLAM
Director, Office of Laboratory Animal Care

University of California, Berkeley
203 NW Animal Facility
Berkeley, CA 94720-7150
