## Supplementary material for "Individually unique, fixed stripe configurations of *Octopus chierchiae* allow for photoidentification in long-term studies": S2 Supporting Information. O. Chierchiae. Breeding and Rearing Methods

*Octopus Chierchiae* Breeding and Rearing Methods

Two male and two female adult *Octopus chierchiae* were obtained from a fully-licensed commercial collector in Nicaragua during 2017 and 2018. We housed them individually in 1-gallon (3.79 liter) glass aquaria lined with gravel, light sand, and bivalve shell fragments [Fig 3A], provided PVC tubes, glass vials, or hollow rocks as shelter, and fed them bait shrimp (*Crangon franciscorum* and *Palaemonetes sp.*) and yellow shore crabs (*Hemigrapsus oregonensis)* ad libitum daily. All aquaria in our laboratory were filled and maintained with artificial seawater (32-36 ppt) at 22-24 ℃.

Beginning in the summer of 2018, we allowed adult *O. chierchiae* (F0 generation) to mate with other wild caught individuals, assumed to be unrelated, by temporarily moving a male into a female’s tank, waiting for up to one hour for copulation to occur, then moving the male back into his tank. Females usually laid eggs within two days of mating, which hatched after about 45 days. Upon hatching, individuals were transferred via a large pipette (turkey baster) into glass jars with screen lids (9 cm tall, 8 cm diameter) containing a variety of substrates including dark sand, light sand, bivalve shell fragments, snail shells, and gravel. Hatchlings [Fig 1B] have a mantle length of around 3.5 mm (Rodaniche, 1984) and immediately begin their demersal lifestyle upon leaving the egg, making it possible to raise them in conventional aquaria.

To avoid issues of cannibalism, territorial disputes, and other aggressive interactions, the animals were reared in individual jars, all within the same 40 gallon tank (151.42 liter) [Fig 3B]. Hatchlings were reared on a diet of primarily marine amphipods and brine shrimp (*Artemia salina*). As they grew and matured, the animals were transitioned to a diet primarily consisting of bait shrimp (*Crangon franciscorum* and *Palaemonetes sp.*) and yellow shore crabs (*Hemigrapsus oregonensis)* and were provided empty snail shells for shelter. Individuals were fed live foods ad libitum when available, and otherwise were offered a piece of shrimp every other day, but displayed an aversion to frozen food. We transferred mature animals of this generation (F1) individually to larger perforated plastic jars in a separate 20 gallon (75.71 liter) tank [Fig 3C]. In order to reduce the risk of genetic strain caused by inbreeding, we selected the least-related male-female pairs and moved them to 1-gallon (3.79 liter) tanks for mating [Fig 3A]. After copulation, we transferred the male back to his plastic jar and provided the female with hollow rocks, PVC pipes, glass vials, and coral fragments in which to lay eggs. Following these methods we were able to successfully close the lifecycle and obtain a generation (F2) bred and raised in captivity.

In our time working with this species, we have observed clutches of up to 32 hatchlings successfully leaving the brood den, and an individual female who, in our care, laid 8 fertile clutches of eggs over the course of two years. As a result, we estimate *O. chierchiae* to be able to produce a maximum of around 240 offspring in its lifetime.
