## Supplementary material for "Individually unique, fixed stripe configurations of *Octopus chierchiae* allow for photoidentification in long-term studies": S4 File. Survey

### Octopus Chierchiae Photo identification Survey

Each of the following questions displays two photos.

Please mark "Match" if you think the photos are of the same individual or "No Match" if you think they are two different animals.

(Note: the image may be taken from different angles and the animals can distort their bodies and colors. Some images have been rotated to show animals in the same orientation)

* Required

1. 1 *

1 point

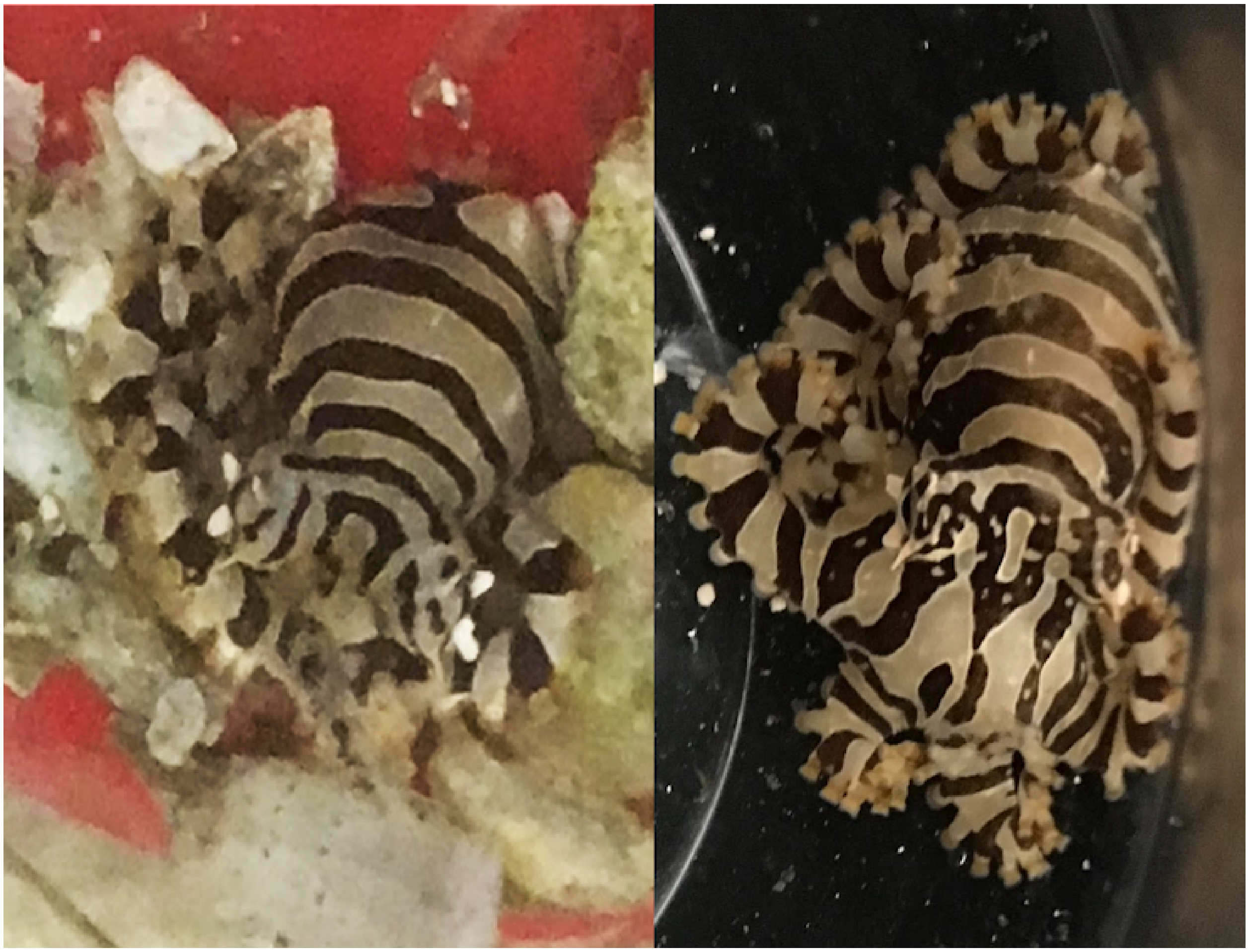

*Mark only one oval.*

- Match
- No Match

2. 2 *

1 point

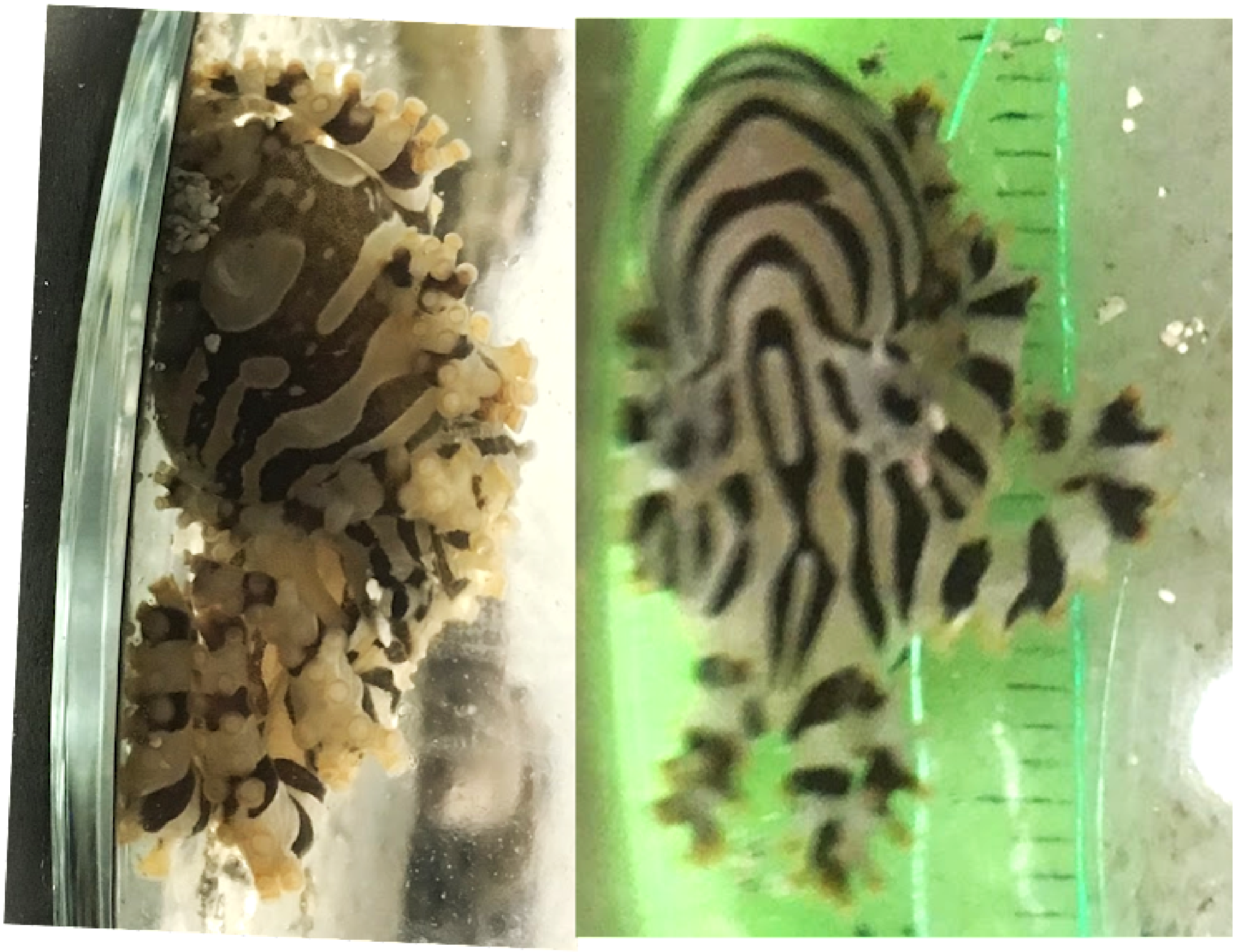

*Mark only one oval.*

- Match
- No Match

3. 3 *

1 point

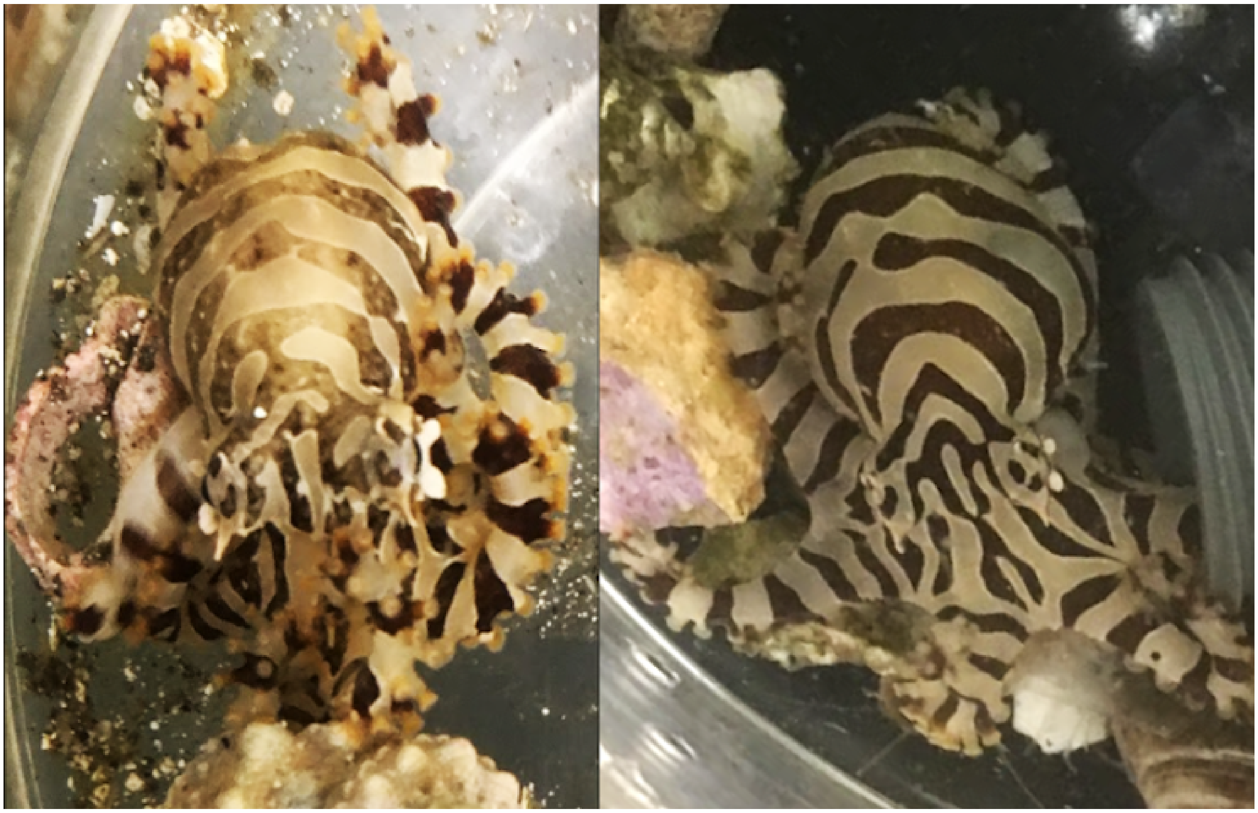

*Mark only one oval.*

- ⬭ Match
- ⬭ No Match

4. 4 *

1 point

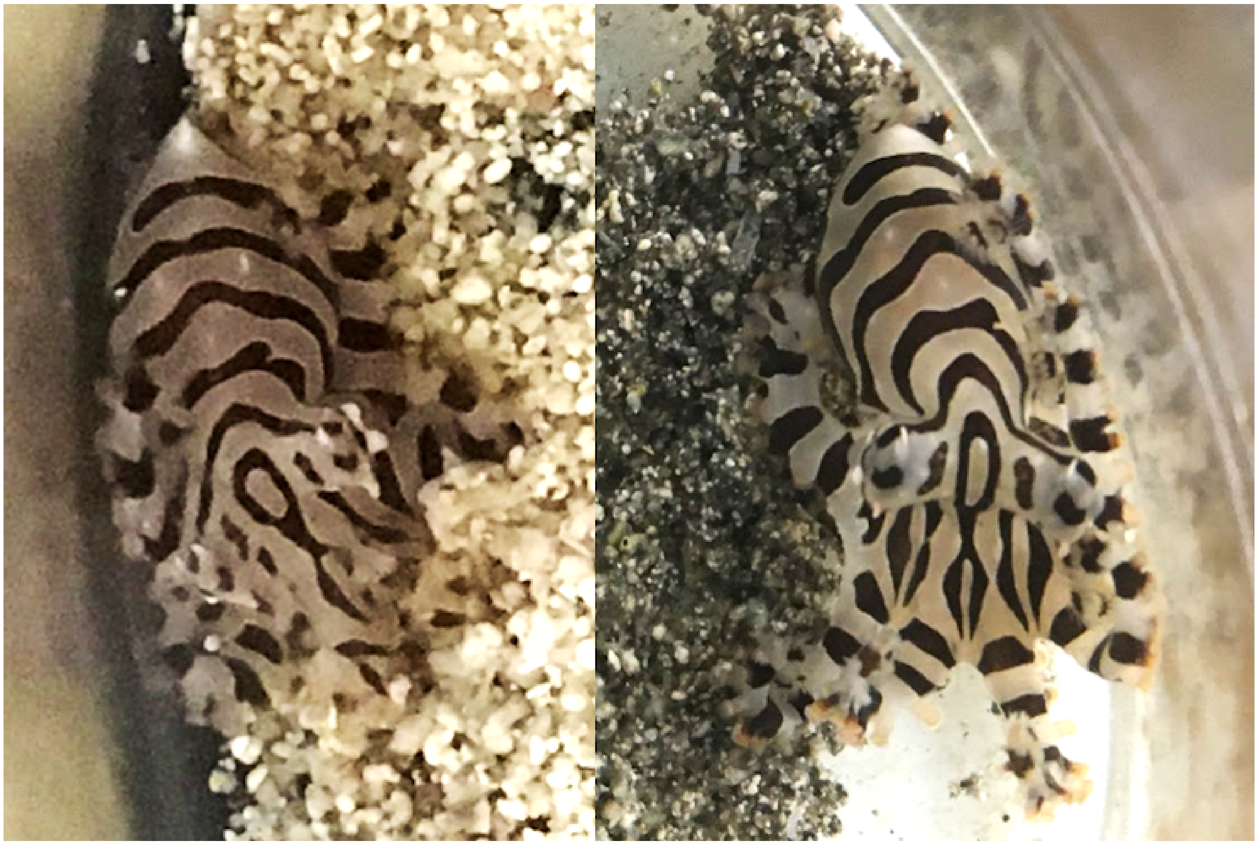

*Mark only one oval.*

- Match
- No Match

5. 5 * 1 point

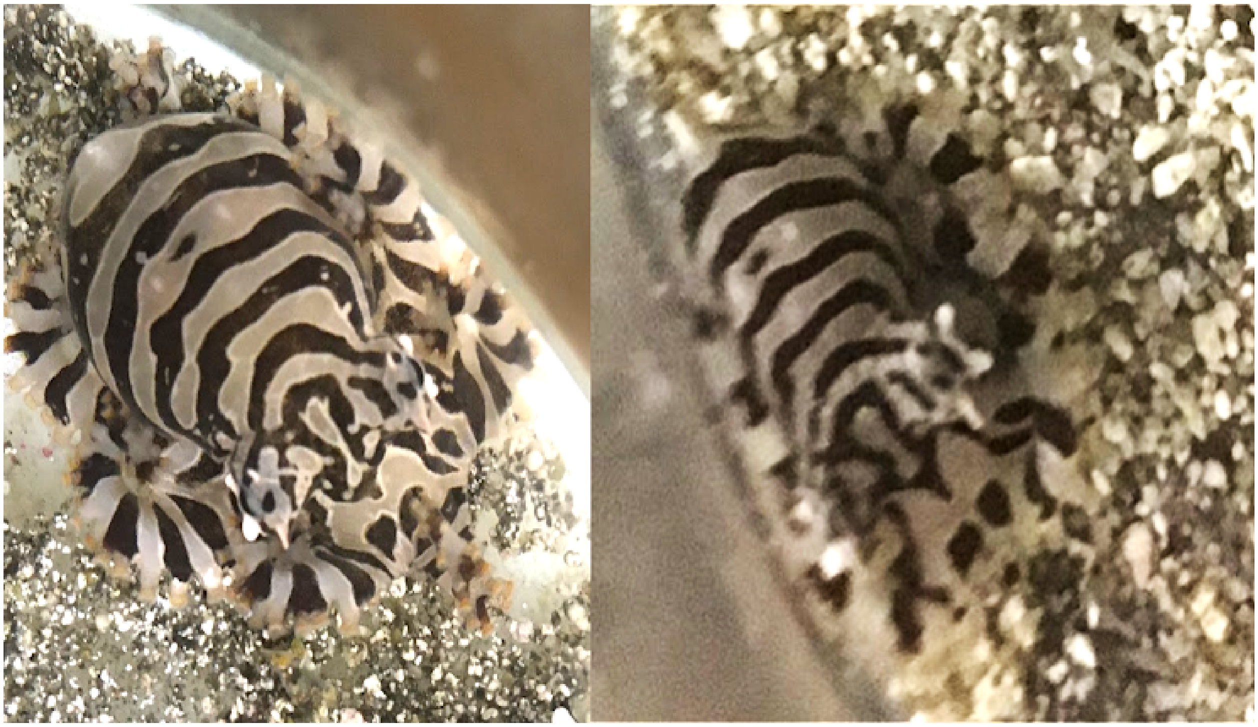

*Mark only one oval.*

- Match
- No Match

6. 6 *

1 point

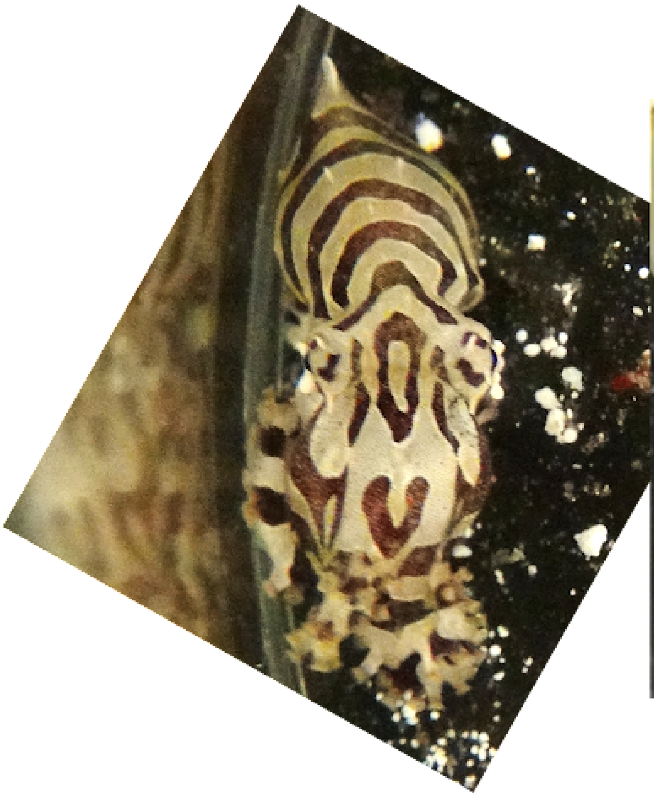

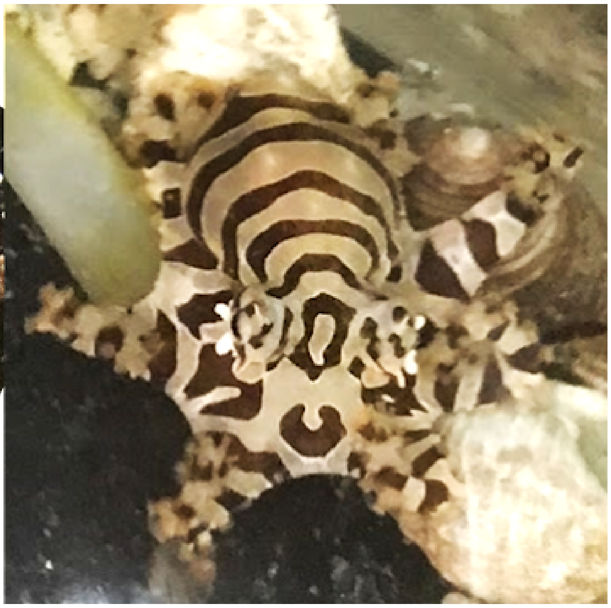

*Mark only one oval.*

- ( ) Match
- ( ) No Match

7. 7 *

1 point

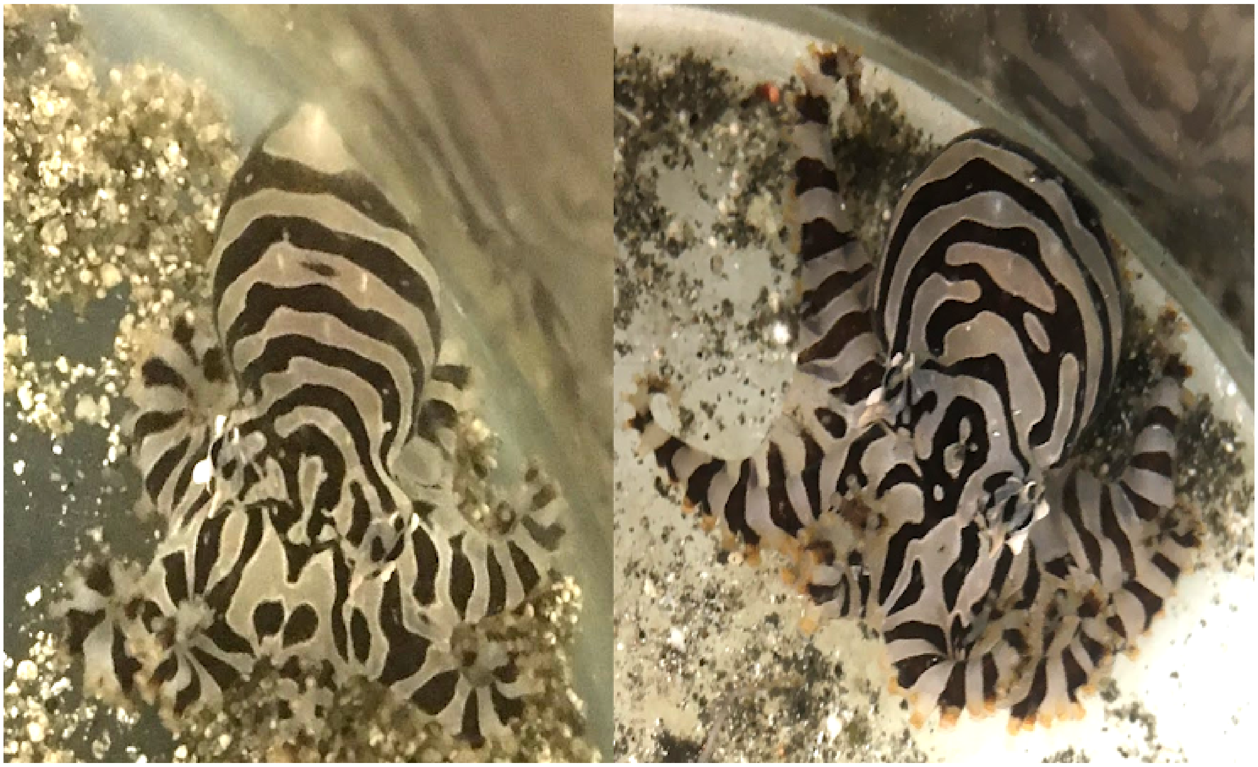

*Mark only one oval.*

- Match
- No Match

8. 8 *

1 point

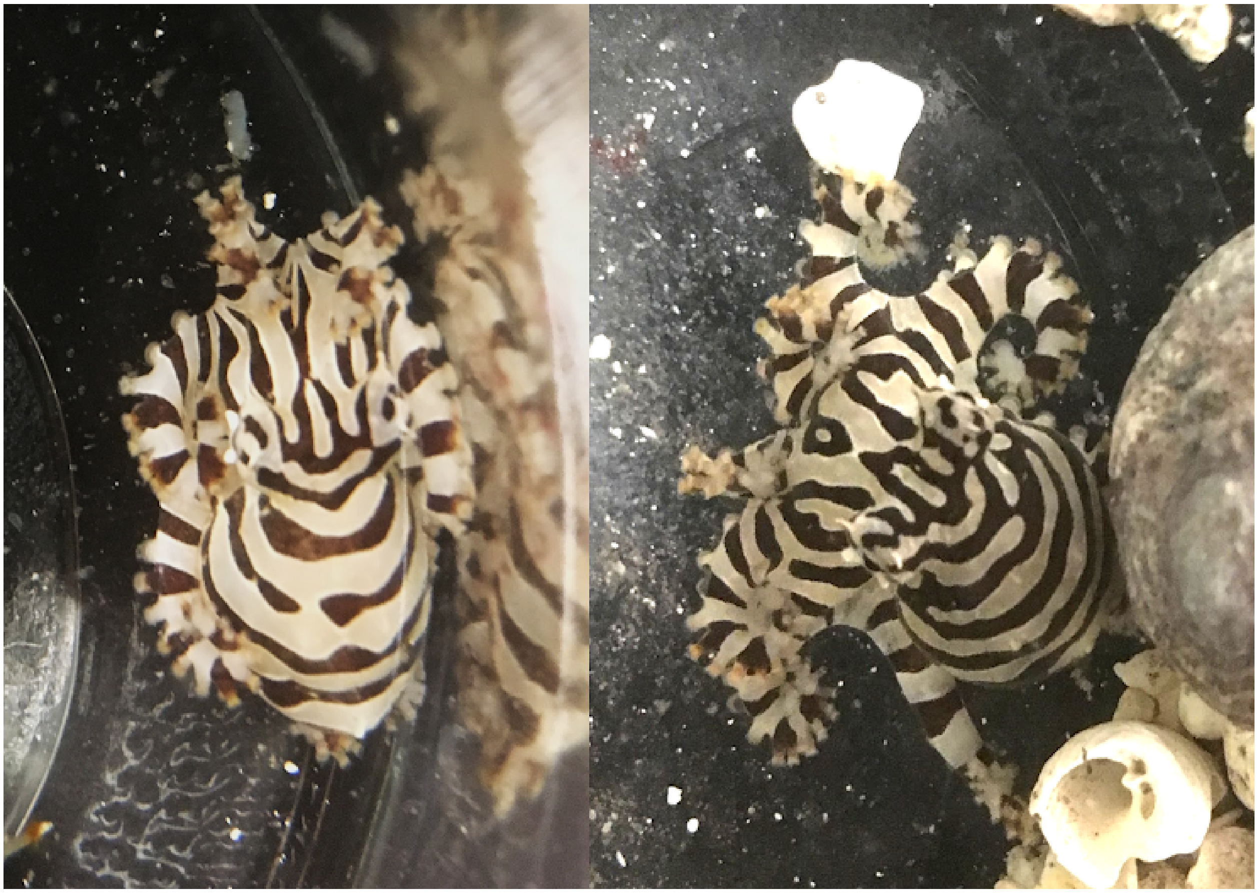

*Mark only one oval.*

- Match
- No Match

9. 9 *

1 point

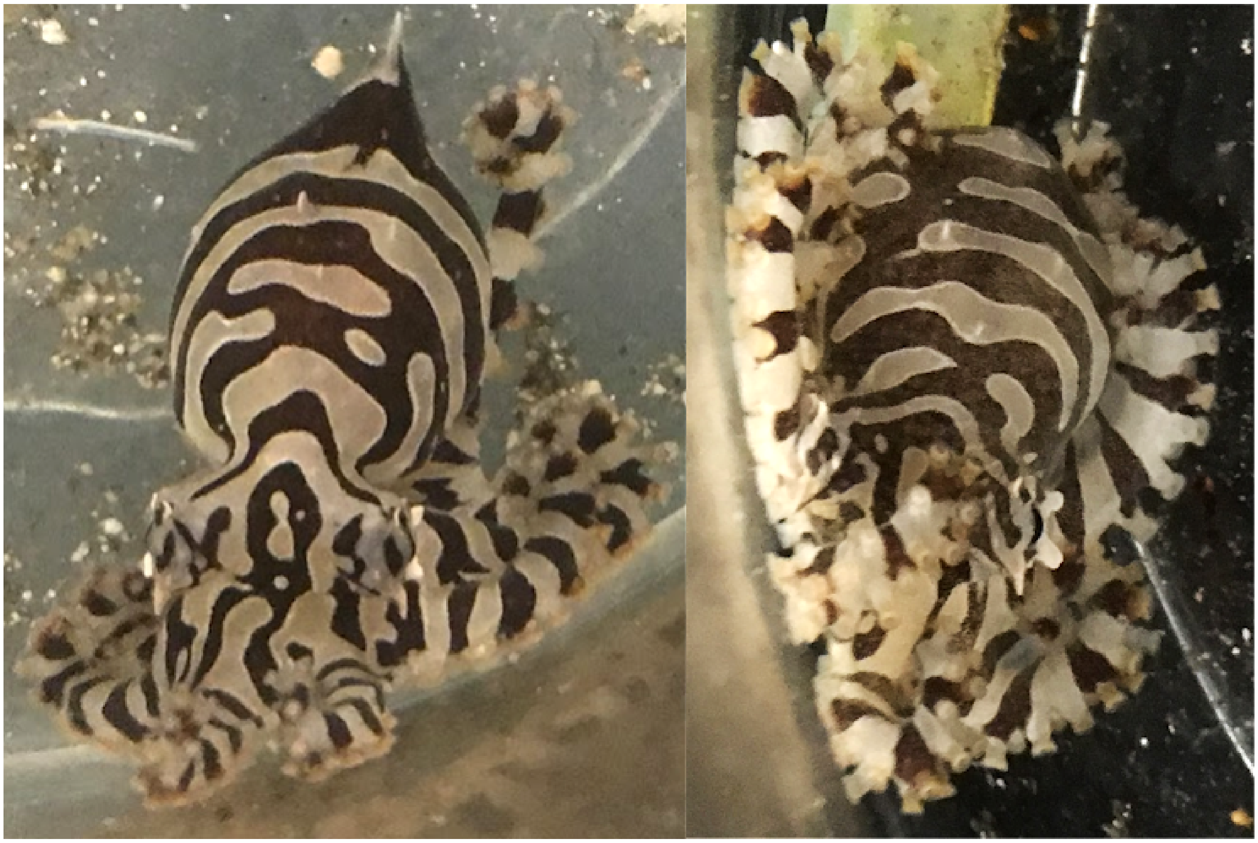

*Mark only one oval.*

- Match
- No Match

10. 10 *

1 point

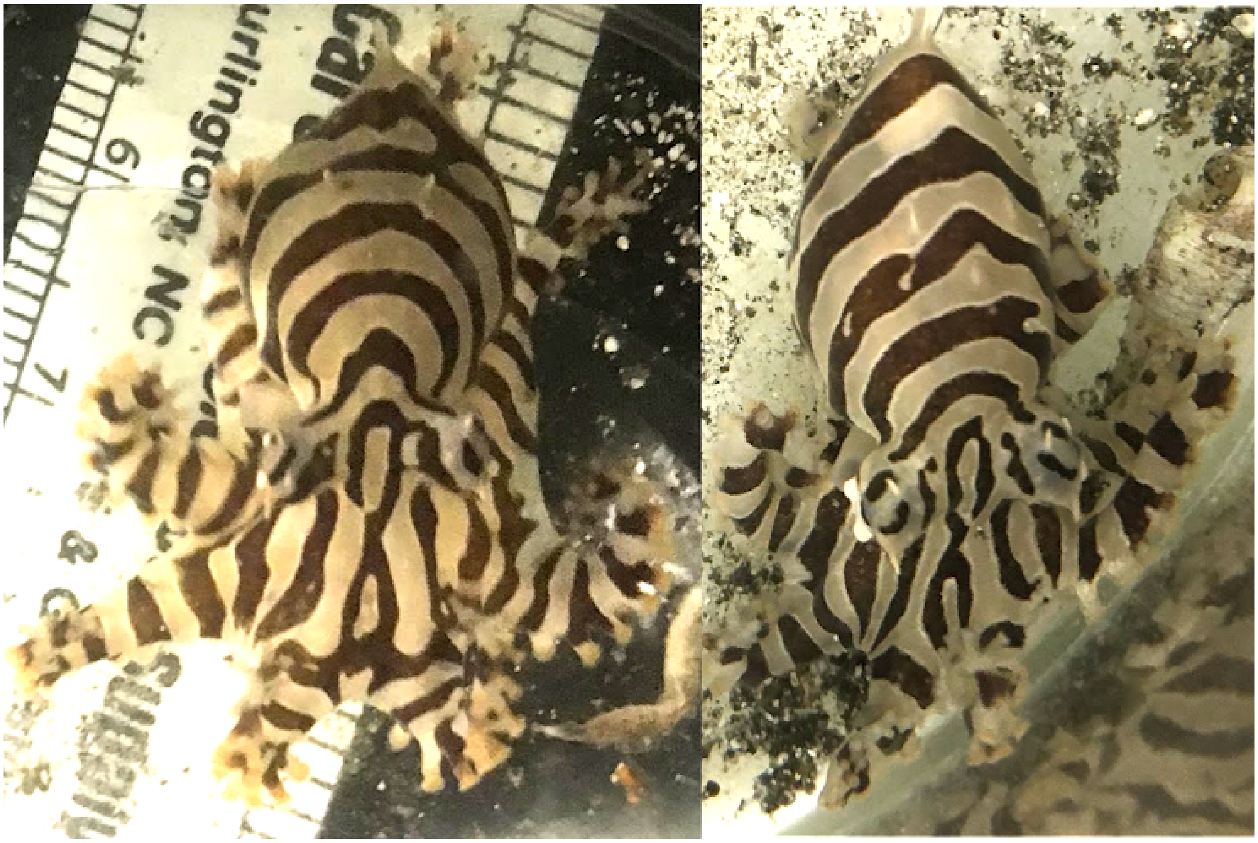

*Mark only one oval.*

- ( ) Match
- ( ) No Match

11. 11 *

1 point

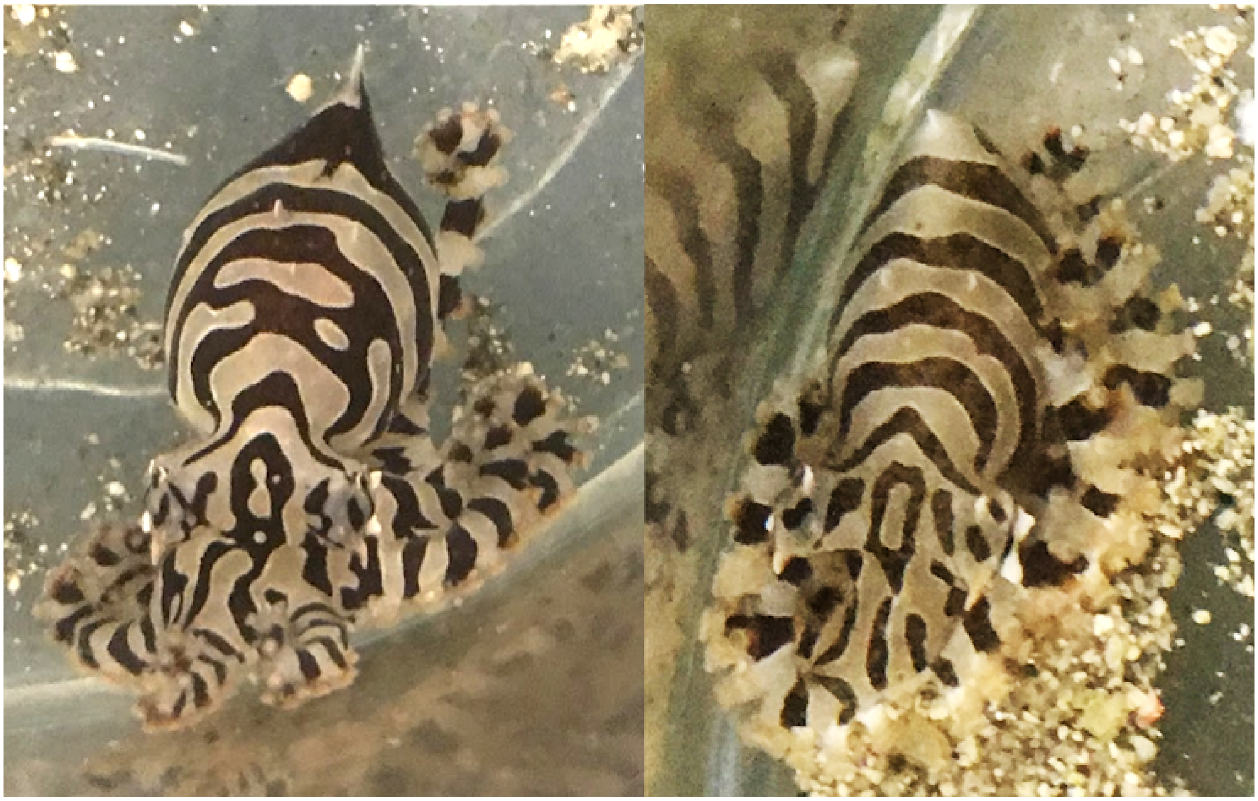

*Mark only one oval.*

- Match
- No Match

12. 12 *

1 point

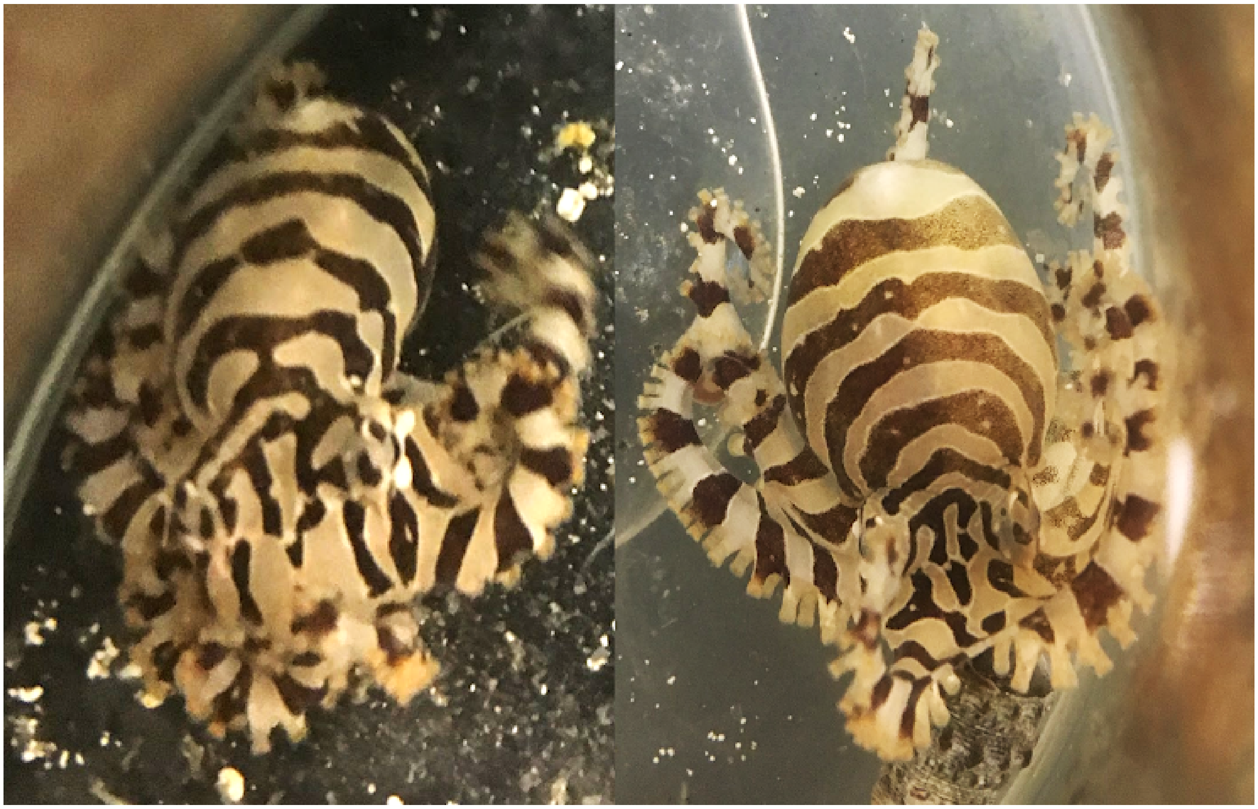

*Mark only one oval.*

- ( ) Match
- ( ) No Match

13. 13 *

1 point

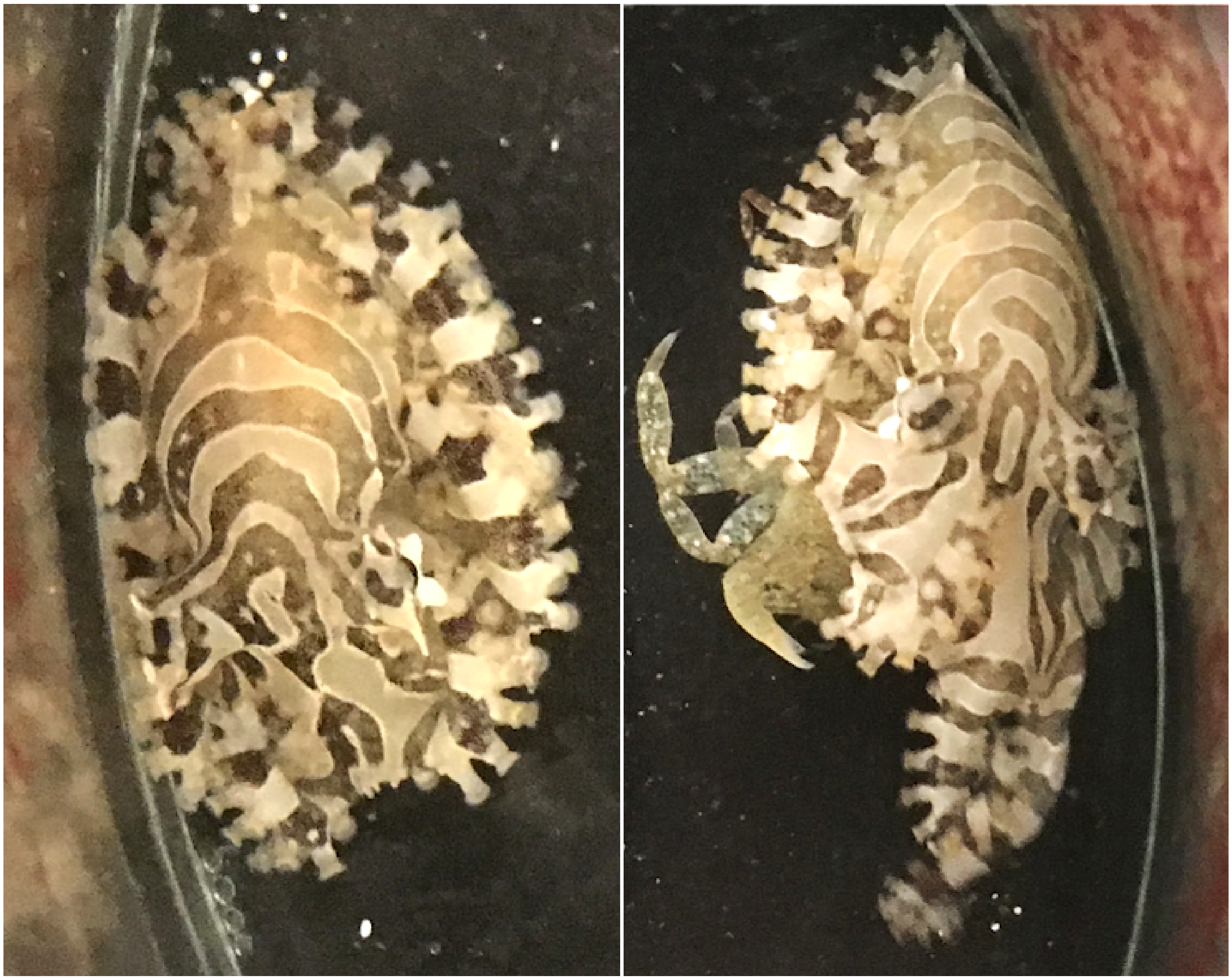

*Mark only one oval.*

- Match
- No Match

14. 14 *

1 point

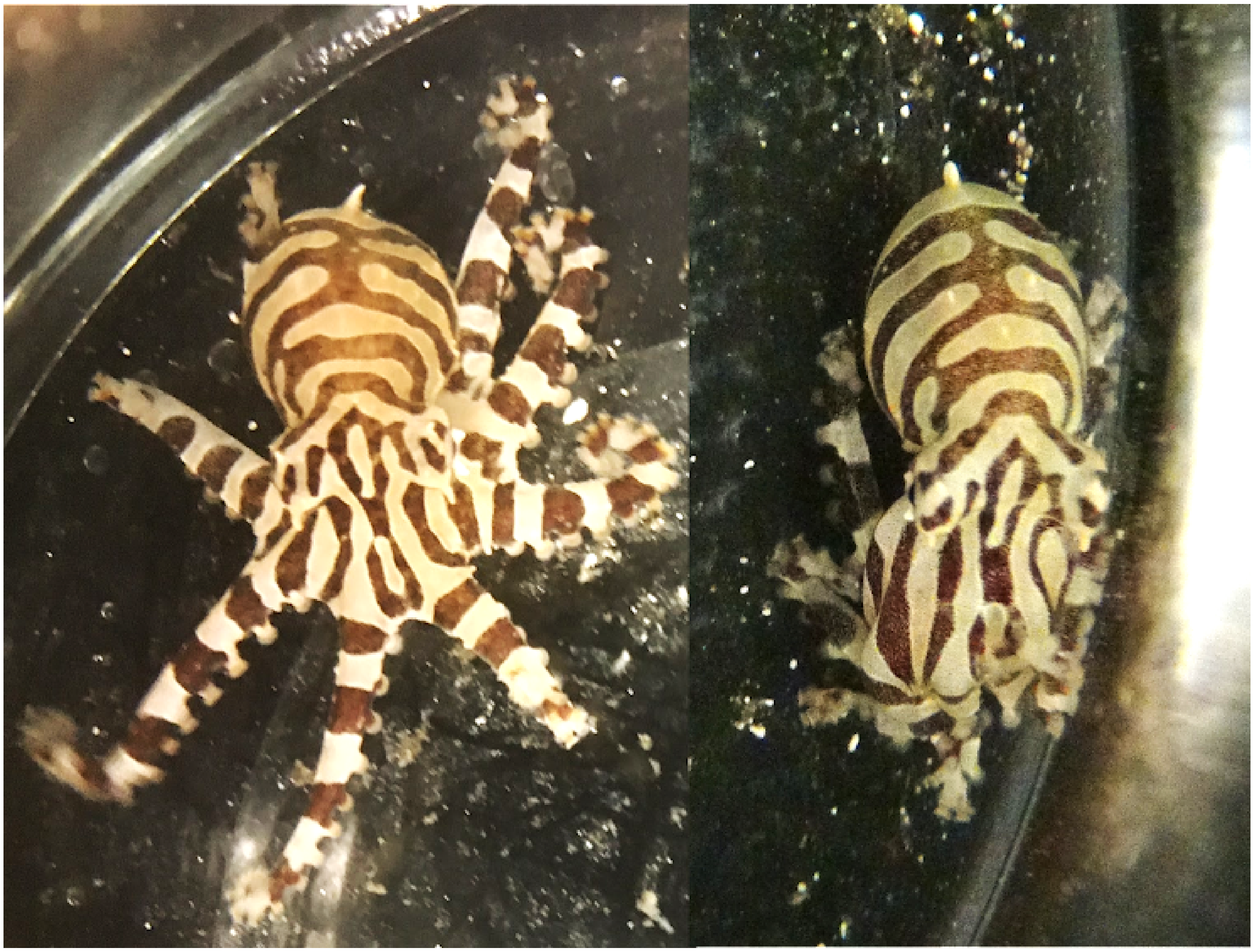

*Mark only one oval.*

- Match
- No Match

15. 15 *

1 point

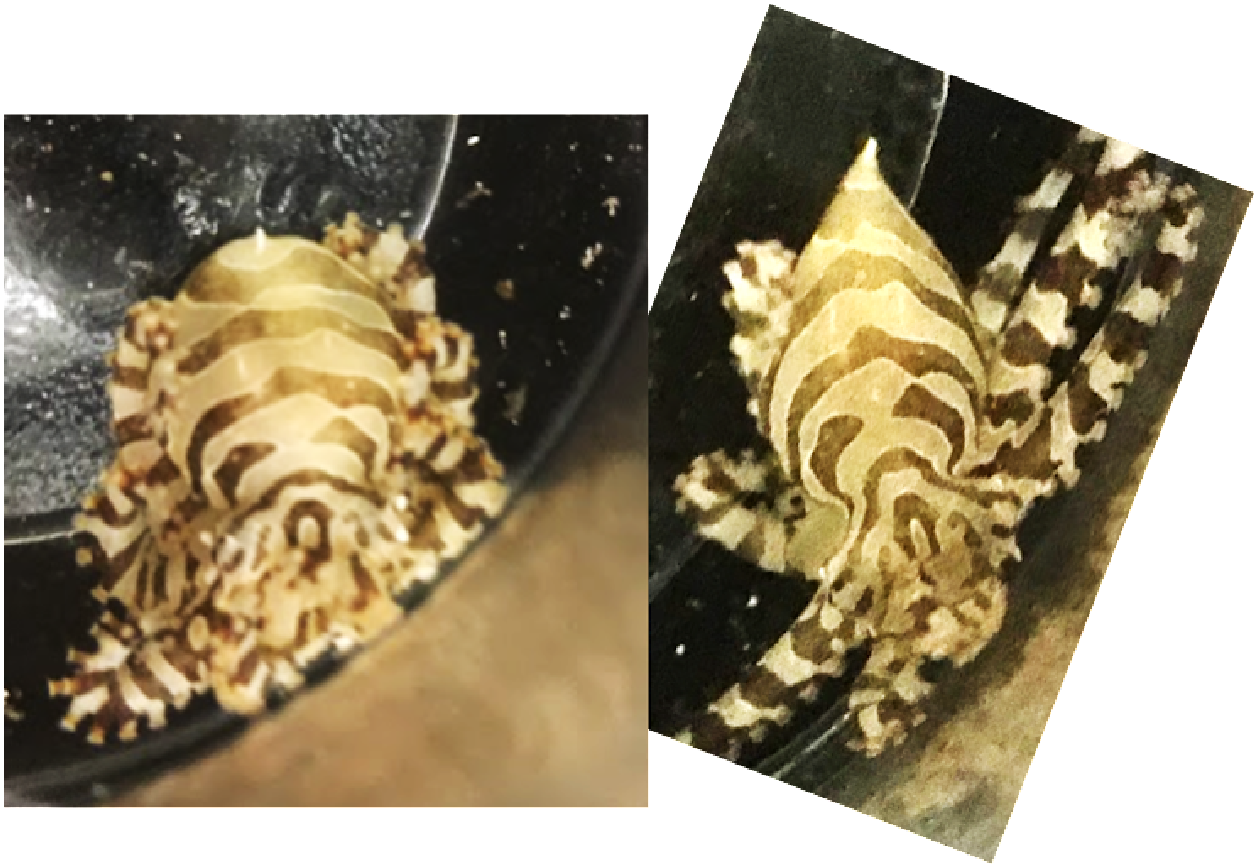

*Mark only one oval.*

- Match
- No Match

16. 16 *

1 point

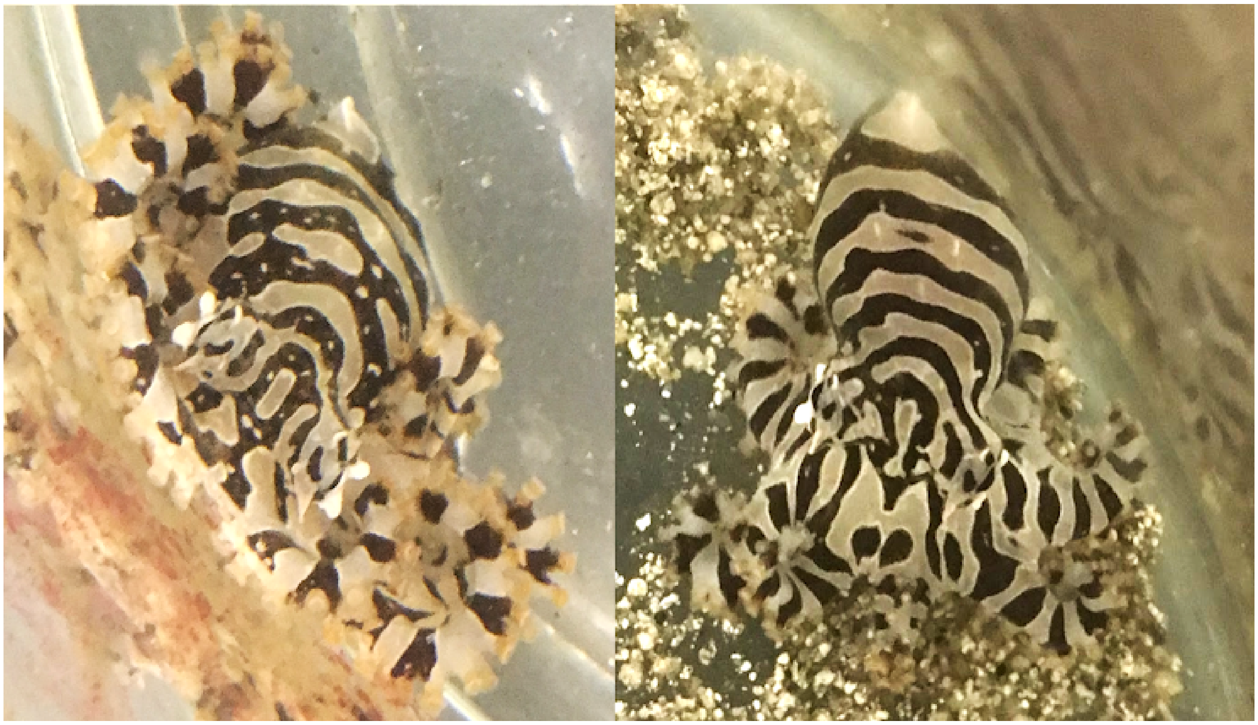

*Mark only one oval.*

- Match
- No Match

17. 17 *

1 point

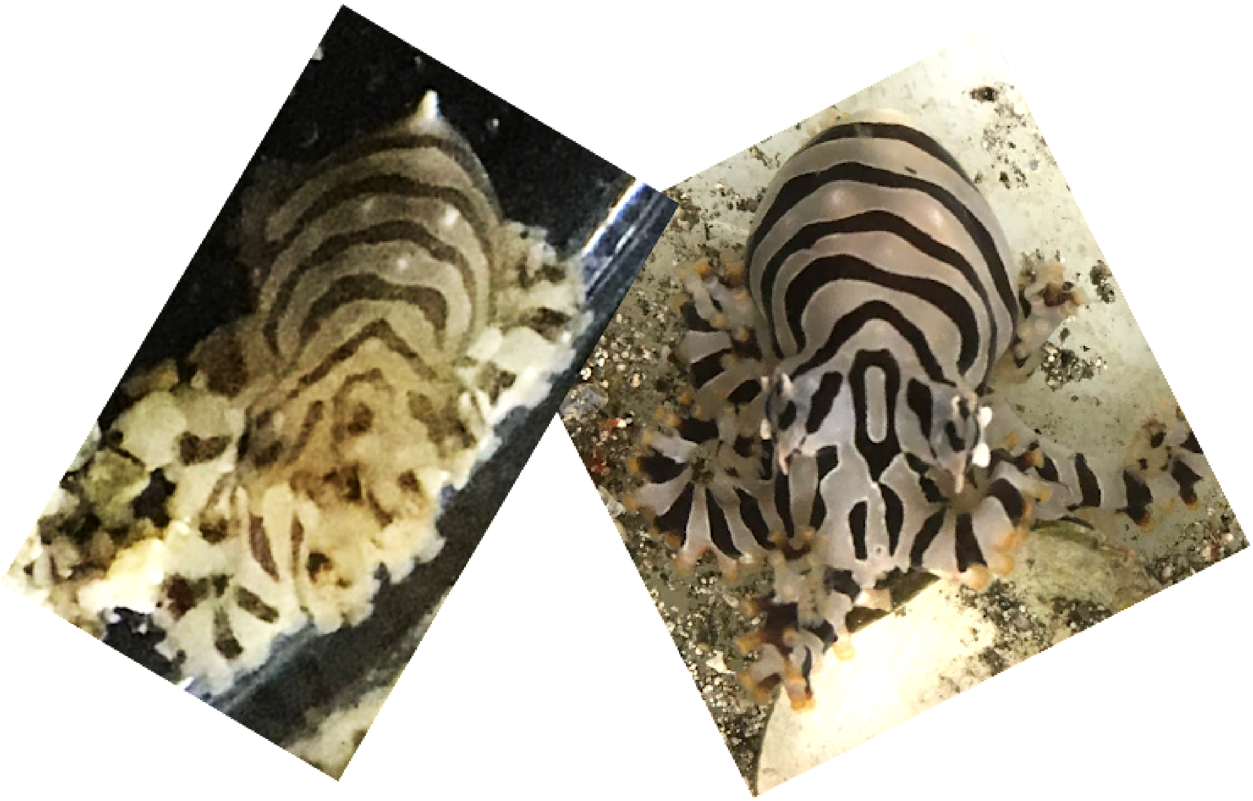

*Mark only one oval.*

- Match
- No Match

18. 18 *

1 point

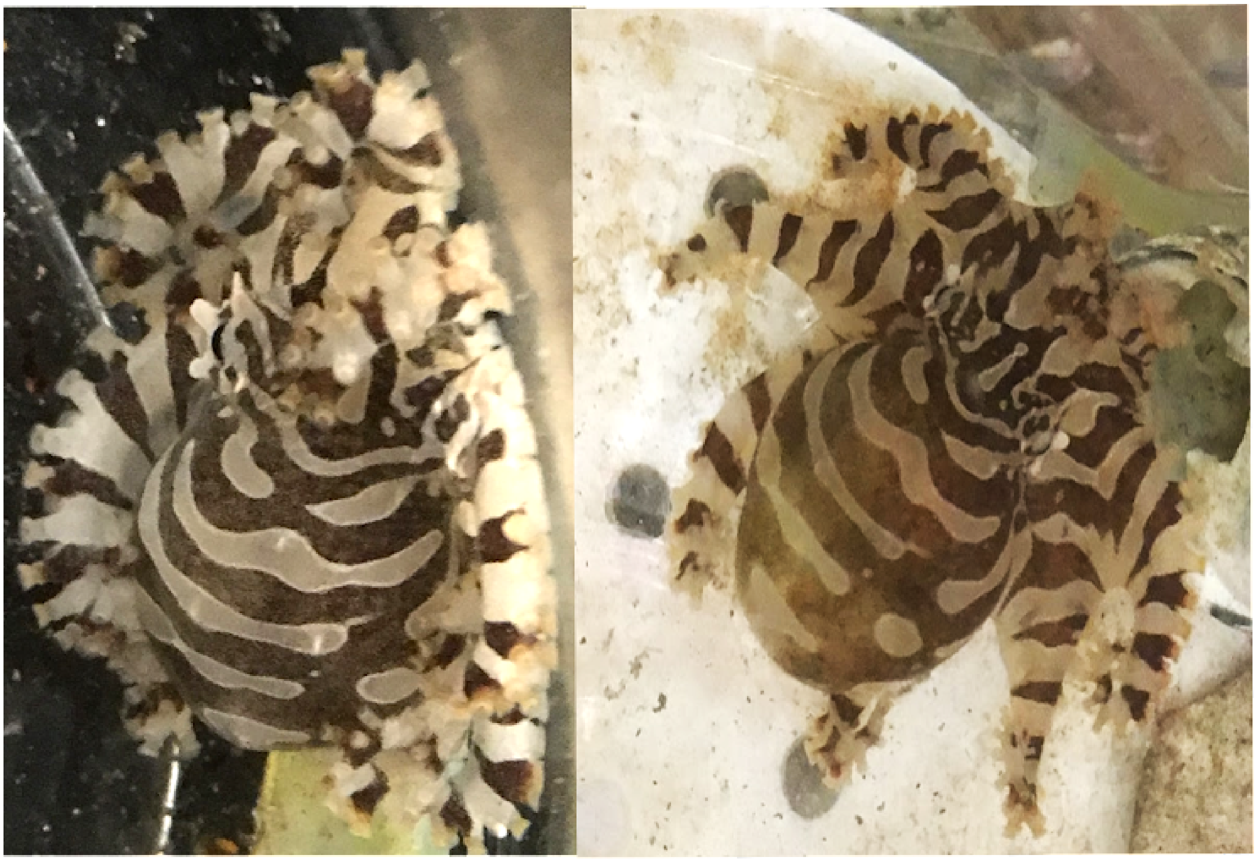

*Mark only one oval.*

- Match
- No Match

19. 19 *

1 point

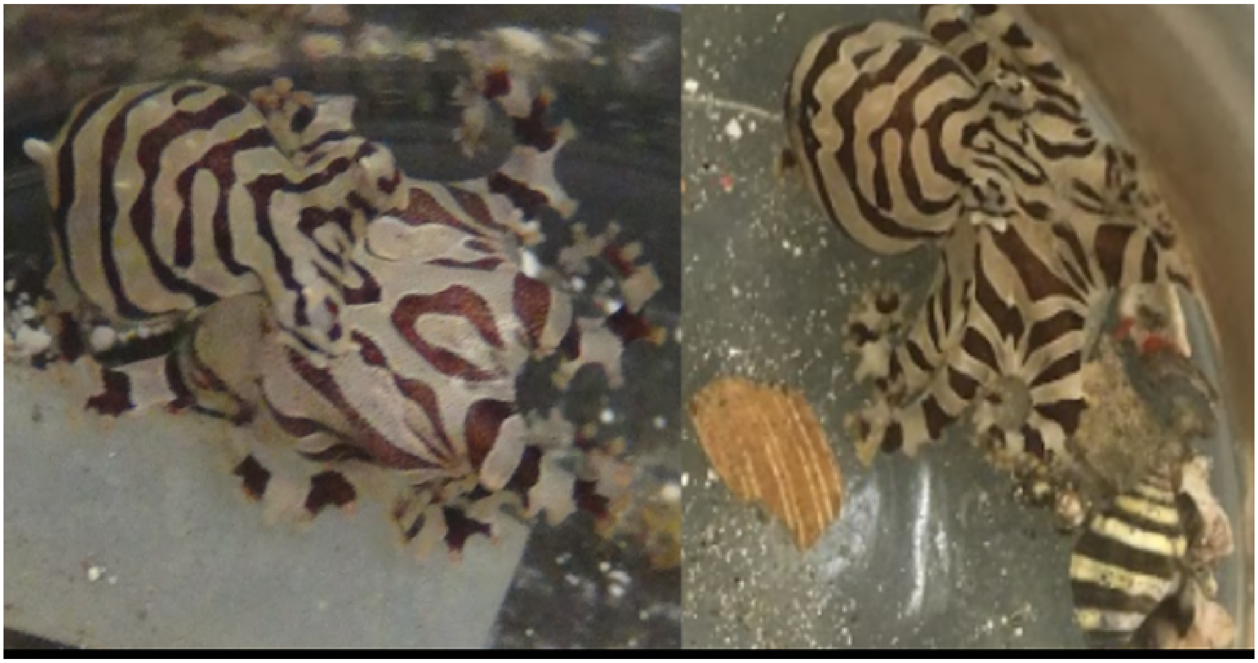

*Mark only one oval.*

- Match
- No Match

20. 20 * 1 point

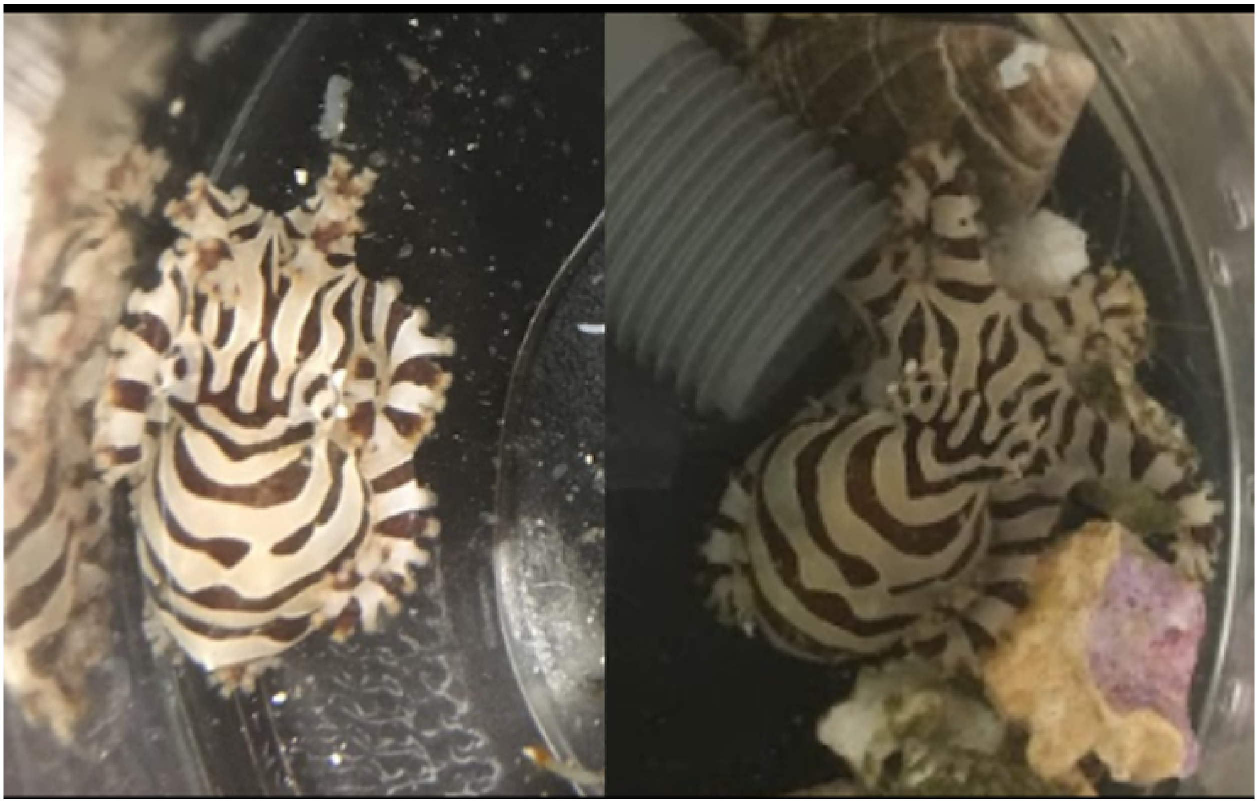

*Mark only one oval.*

- Match
- No Match

This content is neither created nor endorsed by Google.

Google Forms
