## Supplementary figures and images for "Individually unique, fixed stripe configurations of *Octopus chierchiae* allow for photoidentification in long-term studies"

### S8 Figure. Dark Mantle Pattern Component Counts Example

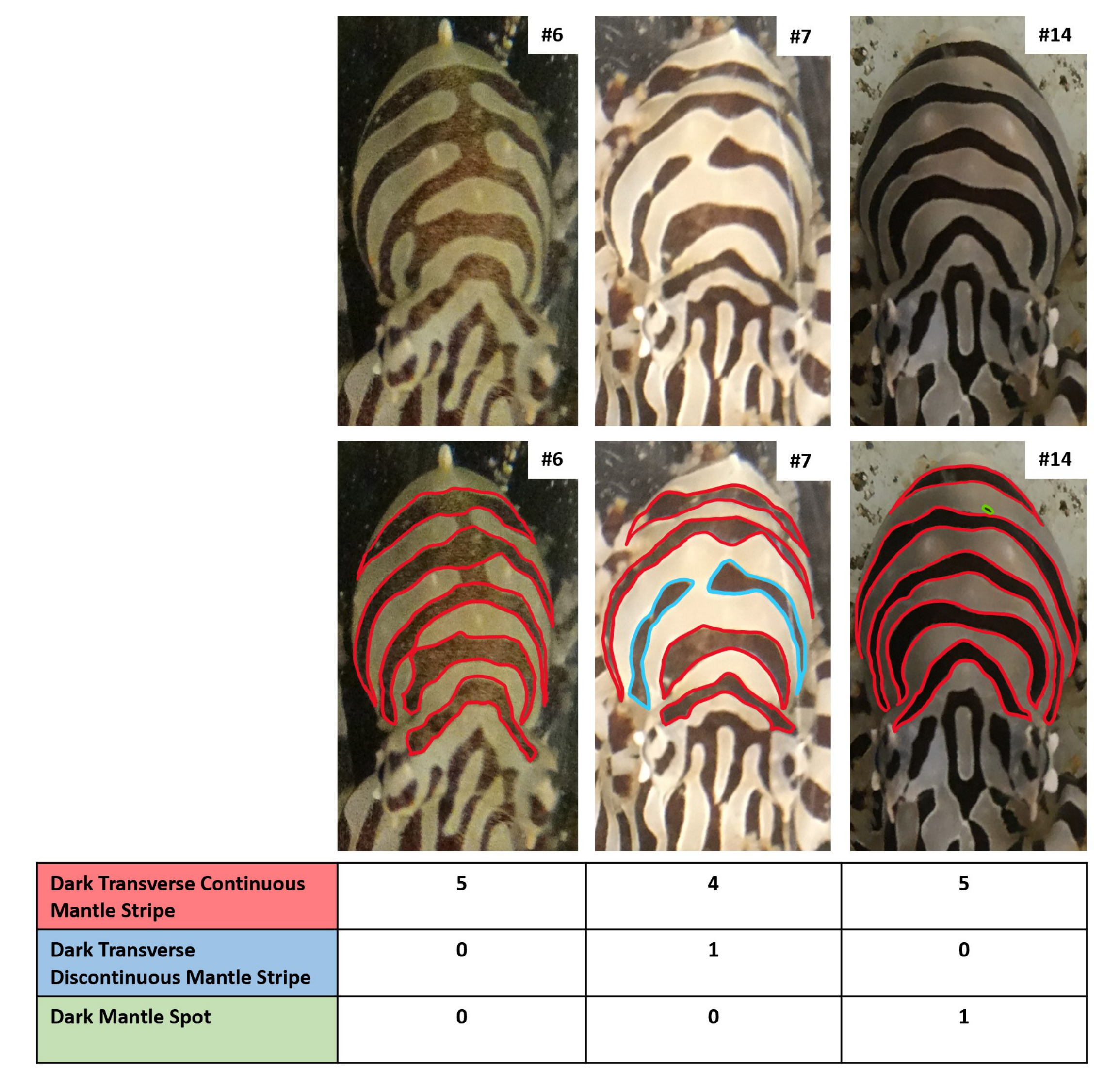
